# The role of hippocampal-vmPFC neural dynamics in building mental representations

**DOI:** 10.1101/2020.04.30.069765

**Authors:** Anna M. Monk, Marshall A. Dalton, Gareth R. Barnes, Eleanor A. Maguire

## Abstract

The hippocampus and ventromedial prefrontal cortex (vmPFC) play key roles in numerous cognitive domains including mind-wandering, episodic memory and imagining the future. Perspectives differ on precisely how they support these diverse functions, but there is general agreement that it involves constructing representations comprised of numerous elements. Visual scenes have been deployed extensively in cognitive neuroscience because they are paradigmatic multi-element stimuli. However, it remains unclear whether scenes, rather than other types of multi-feature stimuli, preferentially engage hippocampus and vmPFC. Here we leveraged the high temporal resolution of magnetoencephalography to test participants as they gradually built scene imagery from three successive auditorily-presented object descriptions and an imagined 3D space. This was contrasted with constructing mental images of non-scene arrays that were composed of three objects and an imagined 2D space. The scene and array stimuli were, therefore, highly matched, and this paradigm permitted a closer examination of step-by-step mental construction than has been undertaken previously. We observed modulation of theta power in our two regions of interest – anterior hippocampus during the initial stage, and in vmPFC during the first two stages, of scene relative to array construction. Moreover, the scene-specific anterior hippocampal activity during the first construction stage was driven by the vmPFC, with mutual entrainment between the two brain regions thereafter. These findings suggest that hippocampal and vmPFC neural activity is especially tuned to scene representations during the earliest stage of their formation, with implications for theories of how these brain areas enable cognitive functions such as episodic memory.

## INTRODUCTION

The hippocampus plays a key role in episodic memory (Scoville & Milner, 1957), spatial navigation (O’Keefe & Dostrovsky, 1971), and a range of other cognitive domains (reviewed in McCormick, Ciaramelli, De Luca, & Maguire, 2018 and Clark & Maguire, 2016). Perspectives differ on precisely how the hippocampus supports these diverse cognitive functions. Nevertheless, there is general agreement that its contribution involves constructing representations comprised of numerous elements (Yonelinas, Ranganath, Ekstrom, & Wiltgen, 2019; Hassabis & Maguire, 2007; Schacter & Addis, 2007; Lee et al., 2005; Cohen & Eichenbaum, 1993). Visual scenes have been deployed extensively to test hippocampal function because they are paradigmatic multi-element stimuli.

We define a scene as a naturalistic three-dimensional (3D) spatially-coherent representation of the world typically populated by objects and viewed from an egocentric perspective (Dalton, Zeidman, McCormick, & Maguire, 2018; Maguire & Mullally, 2013). Whether they are scenes from ongoing experience that are perceived between the interruptions imposed by eye blinks and saccades, or two-dimensional representations (such as photographs) of 3D places, scenes are contexts that you could potentially step into (e.g. a forest) or operate within (e.g. a scene of the desk area in front of you). This stands in contrast to single isolated objects or landmarks on a blank background. These are not scenes and, in fact, are typically deployed as control conditions to compare against scenes (e.g. Hassabis, Kumaran, & Maguire, 2007; Barry, Barnes, Clark, & Maguire, 2019).

Patients with hippocampal damage show scene-related perceptual, imagination and mnemonic impairments (Aly, Ranganath, & Yonelinas, 2013; Mullally, Intraub, & Maguire, 2012; Hassabis, Kumaran, Vann, & Maguire, 2007; Lee et al., 2005), and functional MRI (fMRI) studies have pinpointed the anterior hippocampus in particular as being engaged during scene perception and imagination (Hodgetts, Shine, Lawrence, Downing, & Graham, 2016; Zeidman & Maguire, 2016; Zeidman, Lutti, & Maguire, 2015; Zeidman, Mullally, & Maguire, 2015; Graham, Barense, & Lee, 2010; Hassabis, Kumaran, Vann, & Maguire, 2007). Lesions to another brain region closely connected to the hippocampus (Catani, Dell’Acqua, & Thiebaut de Schotten, 2013; Catani et al., 2012), the ventromedial prefrontal cortex (vmPFC), also adversely affect scene imagination (Bertossi, Aleo, Braghittoni, & Ciaramelli, 2016). Moreover, a recent magnetoencephalography (MEG) study, where participants had 3 seconds to immediately imagine full scenes in response to single cue words (e.g. “jungle”), found theta power decreases in anterior hippocampus and vmPFC, with the latter driving activity in the hippocampus (Barry, Barnes, Clark, & Maguire, 2019).

Nevertheless, it remains unclear whether scenes, rather than other types of multi-feature stimuli, preferentially engage hippocampus and vmPFC. This is important to know because it directly informs theories of how the hippocampus and vmPFC operate and, as a consequence, how they enable cognitive functions such as episodic memory. One account, the scene construction theory, posits that scenes are preferentially processed by the hippocampus (Hassabis & Maguire, 2007; Maguire & Mullally, 2013), with vmPFC driving hippocampal activity (Barry & Maguire, 2019a, 2019b; Ciaramelli, De Luca, Monk, McCormick, & Maguire, 2019; McCormick, Ciaramelli, De Luca, & Maguire, 2018). By contrast, the relational theory argues that it is the associating of multiple elements that requires hippocampal input, irrespective of whether scenes are the outcome of such processing (Cohen & Eichenbaum, 1993; see also Yonelinas, Ranganath, Ekstrom, & Wiltgen, 2019 for a related perspective).

To help adjudicate between these perspectives, Dalton, Zeidman, McCormick, & Maguire (2018) devised an fMRI paradigm which involved gradually building mental representations over three construction stages. This approach was based on previous work that reported three objects and a 3D space seems to be sufficient to generate the mental experience of a scene (Mullally & Maguire, 2013; Summerfield, Hassabis, & Maguire, 2010). Hence, in one condition Dalton, Zeidman, McCormick, & Maguire (2018) had participants build scene imagery from three successive auditorily-presented object descriptions imagined with a 3D space. This was contrasted with constructing images of non-scene arrays that were composed of three objects and a 2D space. The scene and array stimuli were, therefore, highly matched in terms of content and the associative and mental constructive processes they evoked.

Focussing on the medial temporal lobe and averaging across the full imagery construction period, Dalton, Zeidman, McCormick, & Maguire (2018) found that the anterior medial hippocampus was engaged preferentially during the mental construction of scenes compared to arrays during fMRI. Of note, when the imagined 3D and 2D spaces alone (without objects) were examined, neither was associated with increased hippocampal activity. This echoed a previous finding where a 3D space did not provoke engagement of the hippocampus (Zeidman, Mullally, Schwarzkopf, & Maguire, 2012). Consequently, Dalton, Zeidman, McCormick, & Maguire (2018) concluded that it is representations that combine objects with specifically a 3D space that consistently engage the hippocampus.

In the current study we sought to extend previous work to offer novel insights into the construction of mental representations. In the Barry, Barnes, Clark, & Maguire (2019) MEG study, participants imagined a full scene within 3 seconds, and this was compared to a low level baseline (imagining single isolated objects). In the Dalton, Zeidman, McCormick, & Maguire (2018) fMRI study scenes were compared to non-scene arrays, but given the slow nature of the haemodynamic response, it was not feasible to examine the three construction stages separately. By contrast, here we combined the highly matched scene and array construction tasks from Dalton, Zeidman, McCormick, & Maguire (2018) with the high temporal resolution of MEG to study the neural dynamics associated with each of the three construction stages. We could, therefore, provide novel, time-resolved insights into the step-by-step process of scene and non-scene array construction and the specificity of neural responses (if any) to scenes.

We focused this study on two regions of interest, the anterior hippocampus and the vmPFC, given the previous neuropsychological and neuroimaging evidence of their particular importance for scene construction. Moreover, our main interest was in theta. There is a long history linking theta with hippocampal function particularly from rodent studies (reviewed in Colgin, 2016; Kerakas, 2020). Our focus on theta reflects this context, but also the previous MEG study associating theta with immediate scene construction (Barry, Barnes, Clark, & Maguire, 2019). We predicted that theta power would be attenuated relative to baseline. This would be consistent with previous findings of attenuated power associated with the immediate construction of scenes (Barry, Barnes, Clark, & Maguire, 2019; Barry, Tierney et al., 2019) and autobiographical memory recall (McCormick, Barry, Jafarian, Barnes, & Maguire, 2020) using MEG. Moreover, power decreases in frequencies below 30 Hz during memory encoding have been inversely associated with the fMRI BOLD signal (Fellner et al., 2016), and related to enhanced subsequent memory recall (for reviews see Herweg, Solomon, & Kahana, 2020 and Hanslmayr, Staudigl, & Fellner, 2012). Similarly, intracranial EEG studies consistently find decreases in theta during episodic memory (e.g. Fellner et al., 2019; Solomon et al., 2019). Larger decreases observed during scene construction would, therefore, be indicative of increased neural engagement of the hippocampus and vmPFC.

We also had a hypothesis about what would happen across the three construction stages. This prediction took account of previous findings of vmPFC and anterior hippocampal activity during immediate scene construction (where participants had 3 seconds to immediately imagine a full scene; Barry, Barnes, Clark, & Maguire, 2019), and also in the very earliest stage of autobiographical memory retrieval, which also involves scene construction (McCormick, Barry, Jafarian, Barnes, & Maguire, 2020). Concordant with these previous findings, we predicted that anterior hippocampus and vmPFC power changes would be strongest during the first and possibly second stage of scene, but not array, construction.

Different views exist about the nature of the interactions between the hippocampus and cortical areas such as the vmPFC in supporting memory and, we would add, related processes such as scene construction. For instance, some accounts place the hippocampus at the heart of such processing and believe it recruits neocortical regions in the service of this endeavour (Teyler and DiScenna, 1986; Teyler and Rudy, 2007). If this is the case, then the hippocampus should drive activity in vmPFC during the earliest stage of scene construction, and perhaps also during the subseqent stages. This perspective stands in contrast to a model of scene construction that we have previously articulated (McCormick, Ciaramelli, De Luca, & Maguire, 2018; Ciaramelli, De Luca, Monk, McCormick, & Maguire, 2019). Within this architecture, vmPFC initiates the activation of schematic and other knowledge in neocortex that is relevant for a specific scene, while inhibiting elements that are not relevant. This information is conveyed to the hippocampus, which starts to construct the scene. vmPFC then engages in iterations via feedback loops with neocortex and hippocampus to continually update the scene. Hence, using effective connectivity aanalyses for each constuction stage for scenes and arrays we tested three models: (1) hippocampus driving vmPFC; (2) vmPFC driving hipocampus; (3) mutual entrainment between the two regions. We predicted that during the first stage of scene construction the vmPFC would drive the hippocampus. For the subsequent scene construction stages we predicted mutual entrainment between the two areas, reflecting the feedback loops between them.

Our hypotheses aligned with the scene construction theory. However, given the similarity of the scene and array conditions, our paradigm was also capable of revealing if scenes and arrays were treated similarly by the hippocampus, which we believe would be the prediction of relational theorists, given their position that any form of associative processing should recruit this region.

## METHODS

### Participants

Twenty healthy, right-handed participants (13 female; mean age = 25.50 years; SD = 4.70) took part in the experiment. All participants gave written informed consent to participate in accordance with the University College London Research Ethics Committee. All were fluent English speakers with normal vision.

### Stimuli and Task Procedure

During MEG scanning, participants performed two closely matched tasks adapted from Dalton, Zeidman, McCormick, & Maguire (2018). Both tasks involved participants mentally constructing images with their eyes open while looking at a blank grey screen (Figure 1).

**Figure 1.**
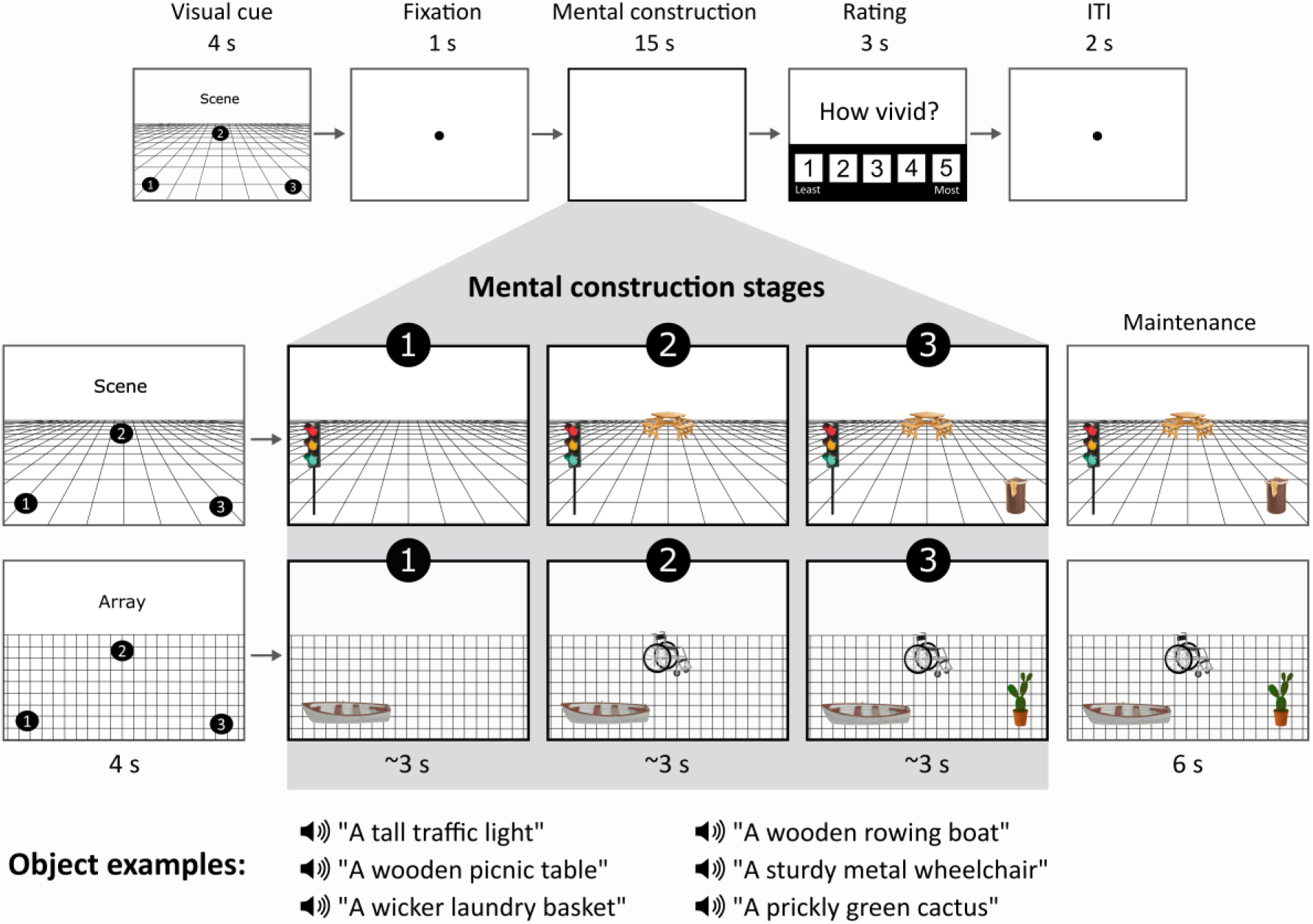
Structure of a trial. The top panel shows the timeline of, in this case, a scene trial. In the middle panels, an example cue configuration is shown on the left, and what participants imagined as they mentally constructed a scene or an array is shown on the right. Of note, during the 3 s construction stages and 6 s maintenance period, participants were looking at a blank screen, and images here only serve as illustrations for the reader. Examples of the auditorily presented object descriptions during each stage are provided in the bottom panel.

For the scene task, participants were asked to first imagine a 3D grid covering approximately the bottom two-thirds of the blank screen. They then heard three object descriptions, presented one at a time, and they imagined each object on the 3D grid, each in a separate location that was specified in advance by the trial cue (Figure 1). The participants were explicitly instructed to link the three objects together and with the 3D space, as they formed their mental representations. Specifically, after the first object description, participants were told to imagine each subsequent object whilst continuing to maintain the image of the grid and previous object(s), without rearranging objects from their original cue-defined positions. By the final object description, the entire mental image created by participants was a simple scene composed of a 3D grid and three objects.

For the array task, participants were asked to first imagine a regular, flat 2D grid covering approximately the bottom two-thirds of the screen. They then heard three object descriptions, presented one at a time, and they imagined each object on the 2D grid, each in a separate location that was specified in advance by the trial cue (Figure 1). The participants were explicitly instructed to link the three objects together and with the 2D space as they formed their mental representations. Specifically, after the first object description, participants were instructed to imagine each subsequent object whilst continuing to maintain the image of the grid and previous object(s), without rearranging objects from their original cue-defined positions. By the final object description, the entire mental image created by participants was a simple array composed of three objects on a flat 2D grid. In this array task it was emphasised to participants they should link the objects and the space together, but not in a way that would create a scene.

For both scene and array tasks, a visual cue determined the three specific locations where participants should imagine each object during a trial. The identity of the objects was not provided at this point, only where they ought to be placed when being imagined while looking at the blank screen. Object descriptions were auditorily presented to the participant during the subsequent mental construction stages (Figure 1). There were four different cue configurations (Figure 2A) which were randomised across the experiment, and the frequency of each was matched across scene and array conditions. We emphasised the importance of following the cue configurations as precisely as possible. This ensured matched eye movements between the scene and array conditions, consistency across participants, and that objects were imagined as separate and non-overlapping, occupying the full extent of the grid. Participants were asked to construct novel mental images of objects based on the descriptions alone, and not to rely on memories associated with the objects described, or to recall specific objects of their acquaintance. Additionally, they were told to imagine only the objects described and not add other elements to a scene or array. All imagery was to remain static and they were required to maintain a fixed viewpoint, as though looking at the image in front of them, rather than imagine moving parts of objects or imagine themselves moving through space.

**Figure 2.**
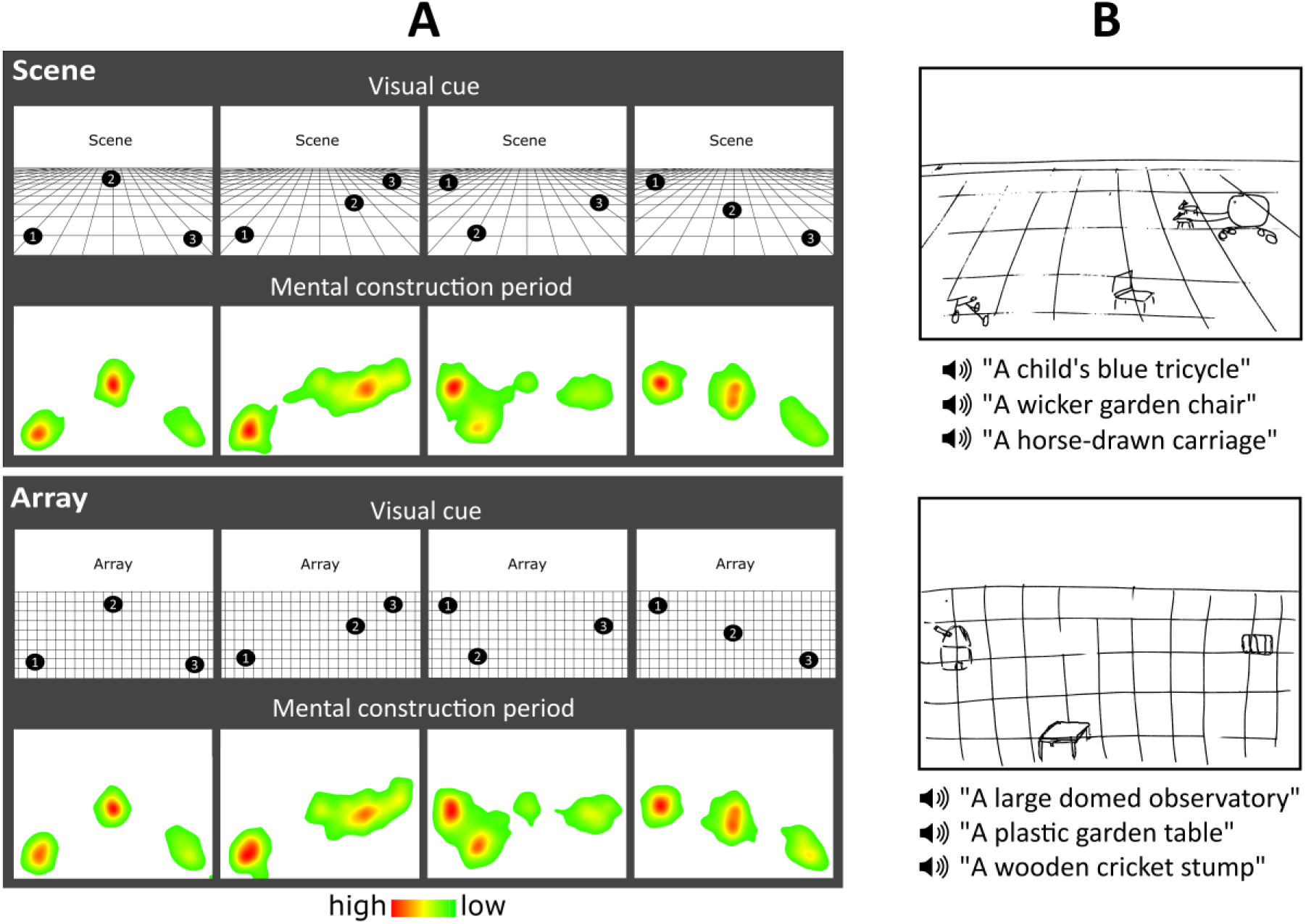
Eye movements and example drawings. (A) Heat maps for each cue configuration for the scene and array tasks show the fixation count during the 9 s mental construction period. Each heat map is an aggregate of fixations on the blank screen across all trials for that cue configuration across all participants with eye tracking data (n=15). Red indicates higher fixation density and green lower fixation density. (B) Pre-scan training drawings for the scene (top) and array (bottom) tasks from an example participant.

The object descriptions were the same as those used by Dalton, Zeidman, McCormick, & Maguire (2018). Each description was heard only once. The objects were rated by a separate group of participants on a range of properties (Dalton, Zeidman, McCormick, & Maguire, 2018), which enabled us to ensure that the scene and array conditions were closely matched in terms of the number and order of presentation of space-defining and space-ambiguous objects (*t*_(71)_ = 0.497, *p* = 0.620) (Mullally & Maguire, 2011; 2013), ratings of object permanence (*t*_(71)_ = 0.178, *p* = 0.859) (Mullally & Maguire, 2011), as well as the number of syllables in the objects descriptions (*t*_(71)_ = 0.327, *p* = 0.745) and utterance duration (*t*_(71)_ = 0.169, *p* = 0.866). Furthermore, all objects were rated as highly imageable, obtaining a score of at least 4 on a scale from 1 (not imageable) to 5 (extremely imageable). Objects in each triplet were not contextually or semantically related to each other – as part of the piloting in the Dalton, Zeidman, McCormick, & Maguire (2018) fMRI study, if a triplet was deemed consistently across participants to contain objects that were related, this triplet was not included in the main experiment. Object triplets were counterbalanced across conditions such that for some participants a triplet was in the scene condition and for others it was in the array condition. Importantly, and to reiterate, during both tasks participants viewed a blank screen, so there was no difference in visual input between conditions.

A third task was also included in the study. Participants heard a backward sequence of three numbers, after which they were instructed to mentally continue counting backwards. The primary function of this task was to provide participants with relief from the effortful imagination required of the scene and array trials. This counting task was not subject to analysis.

To summarise, scene and array tasks were identical except for one manipulation – imagining objects on a 3D grid induced a sense of depth and perspective, resulting in a mental representation of a scene, in contrast to imaging objects on a 2D grid which resulted in a mental representation of a non-scene array. This close matching of the scene with the array tasks meant any differences in neural activity could not simply be attributed to more general mental construction processes. Moreover, the use of MEG meant that we could now examine the step-by-step construction of these representations with millisecond precision.

### Pre-scan Training

Before entering the scanner, participants were trained to become familiar with the task requirements. They completed four practice trials per task on a desktop computer in a darkened room, to liken the experience to that in the scanner environment. After each trial they rated the vividness of the mental imagery on a scale of 1 (not vivid at all) to 5 (extremely vivid). If they gave a rating of 3 or lower on any trial, the instructions and practice items were repeated. A score of 4 or more qualified them to proceed to the MEG experiment. After one of the practice trials per task, participants drew what they were able to imagine (Figure 2B). The main purpose of the drawings was to ascertain whether a participant drew objects corresponding to the auditory descriptions, in the correct cue-specified locations, and on a 3D or 2D grid, indicating whether objects were correctly imagined in a 3D or 2D space. The experimenter checked that all three criteria were met.

### MEG Task Timings

On each trial in the MEG scanner (Figure 1), a visual cue was presented for 4000 ms that contained both the identity of the task (scene or array) and the configuration of locations at which the objects should be imagined on the grid following each auditory object description. Following the cue, participants fixated on the centre of the screen for 1000 ms before the start of the imagination period (∼15 seconds), during which they engaged in mental imagery whilst hearing three auditory object descriptions one after another delivered via MEG-compatible earbuds. Participants were instructed to move their eyes to where they were imagining each object on the screen. Each object description lasted approximately 2000 ms, followed by a silent 1000 ms gap before the presentation of the next object description, so that each object imagination period was 3000 ms in duration. These three construction stages were our main interest, and the focus of the data analyses. After the final object description, there was an additional 6000 ms silent period where participants maintained the image of the fully realised scene or array they had created, which was included in order to avoid an abrupt transition between trials (Figure 1). Participants then rated how vividly they were able to imagine the scene or array on that trial on a scale of 1 (not vivid at all) to 5 (extremely vivid), self-paced with a maximum duration of 3000 ms. An inter-trial interval of 2000 ms preceded the next trial. The Cogent2000 toolbox for Matlab (http://www.vislab.ucl.ac.uk/cogent.php) was used to present stimuli and record responses in the MEG scanner.

A total of 24 trials per task were presented in a pseudorandomised order across four separate blocks – two blocks of twenty and two blocks of nineteen trials. To ensure participants attended to the tasks throughout scanning blocks, we included six catch trials (two per task, including the low level counting backwards task) pseudorandomly presented across all four scanning blocks. During a catch trial, participants were required to press a button when they heard a repeated object description or number within a triplet. The duration of each block was approximately six to seven minutes, an optimal time for participants to remain still and maintain attention during tasks.

### In-scanner Eye Tracking

An Eyelink 1000 Plus (SR Research) eye tracking system with a sampling rate of 2000 Hz was used during MEG scanning to monitor task compliance. The right eye was used for both calibration and data acquisition, to record fixation eye tracking across the full screen. For some participants the calibration was insufficiently accurate, leaving 15 data sets in the eye tracking analyses.

### Post-scan Surprise Memory Test

Following the experiment, participants completed a surprise item memory test for the scene and array tasks, outside of the scanner on a desktop computer. Participants were presented with all individual auditory object descriptions heard during the scanning experiment (72 scene and 72 array objects – 24 trials of each condition comprised of three objects each) and an additional 72 lure object descriptions not previously heard. Auditory descriptions were presented one at a time, and the order of object descriptions was randomised. After hearing each description, participants pressed a button to indicate “yes” if they remembered hearing that object description during the scanning experiment or “no” if they did not. They then rated how confident they felt about their answer on a scale from 1 (low confidence) to 5 (high confidence). Both responses were self-paced, with a maximum of 5000 ms for each. We did not include an associative memory test. This decision was informed by the previous experience of Dalton, Zeidman, McCormick, & Maguire (2018) using the same task during fMRI. They reported that when tested in a post-scan surprise associative memory test, there was no difference between the scene and array conditions, but participants performed at chance. This is not surprising. An associative memory test is very challenging in the context of this paradigm because participants were not explicitly told in advance that memory would be subsequently tested. The tasks were instead designed to be ‘in the moment’ imagination tasks.

### Post-scan Debriefing

Upon completion of the memory test, participants were asked about their experience of performing the scene and array tasks. Ratings completed during debriefing are reported in the Results (also see Table 1).

**Table 1.** Debriefing ratings.

| **Question** | **Rating (1-5 Likert scale)** | **Task** | **Mean ± SD** |
| --- | --- | --- | --- |
| To what degree did you feel your mind was wandering, that you were thinking of things other than the task? | 1 = no mind wandering...<br>5 = always mind wandering | Scene | 1.20 ± 0.41 |
| | | Array | 1.35 ± 0.49 |
| How difficult was it to imagine the objects on the grid? | 1 = easy...5 = very difficult | Scene | 1.75 ± 0.64 |
| | | Array | 1.75 ± 0.72 |
| How difficult was it to follow the cue's instruction of where to imagine the objects on the grid? | 1 = easy...5 = very difficult | Scene | 1.2 ± 0.41 |
| | | Array | 1.15 ± 0.37 |
| Did you have any sense of 3D space during array trials? Please rate the extent to which you felt the image had a sense of depth. | 1 = completely 2D...<br>5 = completely 3D | Scene | 4.3 ± 0.44 |
| | | Array | 1.55 ± 0.51 |
| For the array trials, did you feel that you imagined the objects in a scene at all and tried to repress this? | 1 = always repressed...<br>5 = never had to repress | Array | 4.55 ± 0.60 |

SD = Standard Deviation.

### Behavioural Data Analysis

Ratings for scene and array per-trial vividness during scanning were compared using paired-samples t-tests, as were the ratings from the post-scan debriefing session. Eye tracking comparisons between scene and array tasks were performed using paired-samples t-tests to examine eye movements across the full 9000 ms construction period and, where relevant, using two-way repeated measures ANOVAs which assessed the effects of task (scene, array) and construction stage (first, second, third). For the post-scan surprise memory test, to determine whether memory for object descriptions was above chance for scenes and arrays, one-sample t-tests were performed. The memory test data were also analysed using two-way repeated measures ANOVAs which assessed the effects of task (scene, array) and construction stage (first, second, third). Statistical analyses were performed using SPSS version 25, using a significance threshold of p < 0.05.

### MEG Data Acquisition

MEG data were acquired using a whole-head 275 channel CTF Omega MEG system within a magnetically shielded room, with a sampling rate of 1200 Hz. Participants were scanned in a seated position, with the back of their head resting on the back of the MEG helmet. Head position fiducial coils were attached to the three standard fiducial points (nasion, left and right preauricular) to monitor head position continuously throughout acquisition.

### MEG Data Preprocessing

All continuous MEG data were high-pass filtered at 1 Hz, to eliminate slow drifts in signals from the MEG sensors. A stop-band filter was applied at 48-52 Hz to remove the power line interference, and a stop-band filter of 98-102 Hz to remove its first harmonic. Epochs corresponding to the three scene construction stages and three array construction stages were each defined as 0 to 3000 ms relative to the onset of each object description, while epochs corresponding to the scene and array maintenance periods were defined as 0 to 6000 ms relative to the offset of the third construction stage. Scene and array baseline periods were defined as 1000 to 2000 ms relative to the onset of the inter-trial-interval (ITI) preceding a scene or array trial. This final 1000 ms of the ITI was chosen as it provided the most separation from task-related activity – the on-screen visual cue was not an appropriate baseline since this contained important preparatory information pertinent to the upcoming scene or array to be constructed. Epochs were concatenated across trials for each condition and across scanning blocks.

### MEG Data Analysis

#### MEG Source Reconstruction

Source reconstruction was performed using the DAiSS toolbox (https://github.com/SPM/DAiSS) implemented in SPM12 (www.fil.ion.ucl.ac.uk/spm). The linearly constrained minimum variance (LCMV) beamformer algorithm was used to generate maps of power differences between conditions at each construction stage. This type of beamformer uses weights that linearly map the MEG sensors to source space. Power is estimated at each source location whilst simultaneously minimising interference of contributions from other sources (Van Veen, Van Drongelen, Yuchtman, & Suzuki, 1997). This results in enhanced detection sensitivity of the target source activity. For each participant, a set of filter weights was built based on data from all six construction stages (three for scenes and three for arrays) within the theta band (4-8 Hz) and a 0 to 3000 ms time window relative to object description onset. Coregistration to Montreal Neurological Institute (MNI) space was performed using a 5 mm volumetric grid and was based on nasion, left and right preauricular fiducials. The forward model was computed using a single-shell head model (Nolte, 2003).

At the first level, whole brain theta power was estimated within each construction stage for each condition, creating one weight-normalised image per construction stage per participant. The images were smoothed using a 12 mm Gaussian kernel. We focused this study on two regions of interest, the anterior hippocampus and the vmPFC. ROI analyses were performed using separate bilateral masks covering each of these regions. Masks were created using the AAL atlas in the WFU PickAtlas software (http://fmri.wfubmc.edu/software/pickatlas). The anterior hippocampus was defined from the first slice where the hippocampus can be observed in its most anterior extent until the final slice of the uncus. The posterior hippocampus was defined from the first slice following the uncus until the final slice of observation in its most posterior extent. We included a separate bilateral posterior hippocampal mask for completeness. The vmPFC mask included Brodmann areas 10, 14, 25, and parts of areas 32, 11, 12 and 13. To determine the peak locations of power differences between the scene and array tasks in the anterior hippocampus and vmPFC, at the second level we performed a t-contrast at each of the three construction stages (for example, scene construction stage 1 relative to array construction stage 1). The search volume was then restricted to these regions by performing ROI analyses implemented in SPM12. We used a FWE corrected threshold of p < 0.05 and a spatial extent threshold of 10 voxels for each ROI. Source localised results were visualised using the MNI 152 T1 image in MRIcroGL (https://www.mccauslandcenter.sc.edu/mricrogl). Peaks in the hippocampus and vmPFC provided the seed regions for subsequent effective connectivity analyses. The beamformer analysis was also repeated in the alpha (9-12 Hz) and gamma (31-100 Hz) bands to examine whether effects of task at each stage were confined to the theta range. For completeness, and while not of primary interest, a similar beamformer analysis was performed for the maintenance period (0-6000 ms) that followed the third construction stage of scenes and arrays.

To investigate whether engagement of the anterior hippocampus and vmPFC changed across the three construction stages during each condition, we used the same source extracted theta power obtained with the beamformer for an additional analysis. Using the previously defined masks (anterior hippocampus, vmPFC and posterior hippocampus), we extracted ROI-specific power values from each of the smoothed first level images. Power was subsequently averaged across each ROI, to produce an average for each participant at each of the six construction stages. The same source reconstruction protocol was performed for a pre-trial fixation baseline time-window, which was then subtracted from each task stage to ascertain whether theta power originating from each ROI during task engagement corresponded to an increase or attenuation of power. Data were then entered into two-way repeated measures ANOVAs implemented in SPSS25, allowing us to evaluate changes in power between scene and array tasks and across construction stages (task × stage). Where the assumption of sphericity was violated using the Mauchly test, Greenhouse-Geisser corrected degrees of freedom were reported. Paired-sample t-tests were performed for each ROI in cases of a significant main effect or interaction, using a conservative significance level of p < 0.01.

#### Effective Connectivity

To assess how the hippocampus and vmPFC interacted during each of the three construction stages for both scenes and arrays, we used Dynamic Causal Modelling (DCM) for cross spectral densities (Moran et al., 2009), which measured the magnitude of cross spectra between the ROIs. This connectivity approach permitted the comparison of biologically plausible models representing a priori hypotheses of the directed influence one region has over another, as well as mutual entrainment between regions (Kahan & Foltynie, 2013; Friston, 2009).

Electrophysiological DCMs use neural mass models that summarise the activity within a specified cluster of neurons. We employed a convolution-based local field potential biophysical model, used for EEG/MEG data (Barry, Barnes, Clark, & Maguire, 2019; Moran, Pinotsis, & Friston, 2013; Moran et al., 2009). Intrinsic connectivity between different cell populations within a region are estimated. Extrinsic afferent inputs are categorised as forward, backward or lateral depending on which subpopulations these afferents project to (Felleman & Van Essen, 1991). Forward connections project to spiny stellate neurons in middle cortical layers, backward connections project to both excitatory pyramidal neurons and inhibitory interneurons, and lateral connections project to all cell populations.

Random-effects (RFX) Bayesian model selection (BMS) was then used to compare the evidence for different competing models that varied according to which connections were modulated by the experimental condition (Stephan, Penny, Daunizeau, Moran, & Friston, 2009). The RFX procedure does not assume the optimal model is the same across all participants, making it more accurate and robust against outliers than fixed-effects, since variability in brain activity across participants is to be expected when studying a complex cognitive process (Stephan et al., 2010). For each construction stage, we tested three models corresponding to the three hypotheses regarding hippocampal-vmPFC connectivity: (1) hippocampus drives vmPFC, (2) vmPFC drives hippocampus, and (3) hippocampus and vmPFC are mutually entrained. When applied to our data, we determined the winning model to be the one with the greatest exceedance probability, which provides a measure of how likely one model is, compared to all other models across the group of participants as a whole. Evidence for each model was a balance between the goodness-of-fit of the model with how parsimonious an explanation it provided, using the minimum number of parameters possible.

## RESULTS

The goal of this study was to examine whether the anterior hippocampus and vmPFC supported step-by-step scene construction whilst using the array condition to control for neural responses associated with objects, and general associative and constructive processing. We first examined the behavioural data to assess whether the participants adhered to the task instructions and to ascertain if there were any differences between the scene and array tasks that might have influenced the neural activity.

### Behavioural Results

#### Did Participants Engage in the Scene and Array Tasks and Adhere to Instructions?

Several methods were used to assess whether participants paid attention during scene and array trials, engaged in mental imagery, and complied with task-specific instructions throughout the experiment.

#### Pre-scan training

After performing scene and array practice trials during the pre-scan training, participants drew what they imagined on each trial, to ensure they could closely follow the task instructions and cue configurations. Scene and array drawings demonstrated participants formed the appropriate mental representations, related the objects and the space together in a coherent way, and were able to mentally place the objects in the locations indicated by the cue configurations (see examples in Figure 2B).

#### Catch trials

Catch trials were included in each scanning block whereby participants were instructed to press a button whenever they heard a repeated object description (or number) within a trial. On average 98.33% (SD = 0.31) of catch trials were correctly identified, suggesting participants remained attentive throughout the experiment.

#### In-scanner eye tracking

Heat maps of the spatial pattern of fixations for the cue configurations confirmed not only remarkably close adherence to the instructions trial-by-trial, but also consistency across participants (Figure 2A). The eye tracking data were also examined for any differences between the scene and array tasks over the full 9000ms construction period, of which none were evident – fixation count (*t*_(14)_ = 0.239, *p* = 0.814), fixation duration (*t*_(14)_ = 0.559, *p* = 0.585), saccade count (*t*_(14)_ = 0.322, *p* = 0.752), saccade amplitude (*t*_(14)_ = 0.752, *p* = 0.465). We also examined the two fixation variables for any effects of task (scene, array) and construction stage (first, second, third). For both fixation count and fixation duration, there was no significant effect of task (fixation count: *F*_(1, 14)_ = 1.441, *p* = 0.250; fixation duration: *F*_(1, 14)_ = 2.964, *p* = 0.107) or construction stage (fixation count: *F*_(2, 28)_ = 0.372, *p* = 0.693; fixation duration: *F*_(2, 28)_ = 0.650, *p* = 0.530), and there was no significant task x construction stage interaction (fixation count: *F*_(2, 28)_ = 2.394, *p* = 0.110; fixation duration: *F*_(2, 28)_ = 0.356, *p* = 0.704). Eye movements were therefore closely matched across all construction stages, and there was no significant difference between the scene and array tasks.

### Did Vividness of Mental Imagery Differ Between Scene and Array Tasks?

Participants rated the vividness of mental imagery immediately after each scene and array trial in the scanner on a scale of 1 (not vivid at all) to 5 (extremely vivid). On average, both scene (mean = 4.15, SD = 0.43) and array (mean = 4.08, SD = 0.49) trials were rated high on vividness, and there was no significant difference between the tasks (*t*_(19)_ = 0.873, *p* = 0.394).

### Were There Any Other Subjective Differences in Mental Imagery Between the Scene and Array Tasks?

After the experiment was completed, participants reported on their subjective experience of performing scene and array tasks in a debriefing session (questions and summary data are shown in Table 1). Mind-wandering was reported as very low for both scene and array tasks, and did not differ significantly between the two conditions (*t*_(19)_ = 1.831, *p* = 0.083). Mental imagery was performed with ease in both conditions (*t*_(19)_ = 0.000, *p* = 1.000). Participants also had no difficulty following the cue configuration instructions for either condition (*t*_(19)_ = 1.000, *p* = 0.330). As expected, scene trials were perceived as significantly more 3D than array trials, while arrays were rated as more 2D (*t*_(19)_ = 17.168, *p* < 0.0001). Participants also reported that they rarely needed to suppress scene-like mental imagery during array trials.

### Did Memory Performance Differ Between Scene and Array Tasks?

After scanning, participants immediately engaged in a surprise item memory test. They performed above chance at recognising individual object stimuli from scene trials (mean = 84.31%, SD = 9.63; *t*_(19)_ = 39.135, *p* < 0.0001), array trials (mean = 85.42%, SD = 9.43; *t*_(19)_ = 40.482, *p* < 0.0001) and identifying novel items (mean = 87.54%, SD = 7.48; *t*_(19)_ = 52.289, *p* < 0.0001). Participants, therefore, paid sufficient attention during both tasks to successfully encode a large number of stimuli, even though they were never instructed to memorise. A repeated-measures ANOVA revealed no significant effect of task (*F*_(1, 19)_ = 0.456, *p* = 0.508), construction stage (*F*_(1.487, 28.262)_ = 1.480, *p* = 0.240), and no task x construction stage interaction (*F*_(1.554, 29.526)_ = 0.739, *p* = 0.484).

### Behavioural Results Summary

Having determined across a range of behavioural measures that the participants adhered to the task instructions, formed the appropriate mental images, with no differences in terms of key variables such as eye movements, vividness and memory performance between the scene and array tasks, we next examined the neural data.

## MEG Results

Our ROIs were the anterior hippocampus and vmPFC, and our primary question was whether they would be preferentially engaged during the step-by-step construction of scenes.

### Source Space Power Differences for Each Construction Stage Separately

We first compared the scene and array tasks during each of the three construction stages. For the first stage we observed a significant attenuation of theta power for the scene relative to the array task in the left anterior hippocampus (peak x, y, z = −24, −20, −16; Figure 3A), left vmPFC (peak in BA32, −12, 40, −14; Figure 3A) and right vmPFC (peak in BA11, 16, 26, −18). The contrast of scene and array conditions during the second construction stage revealed an attenuation of theta activity for scenes in the vmPFC, extending bilaterally with an overall peak (−4, 26, −4) and subpeak (−12, 32, −18) in the left hemisphere (Figure 3A). No significant differences in anterior hippocampus or vmPFC were apparent between the scene and array tasks during the third construction stage, or during the ensuing maintenance period. For completeness, no task-related theta power changes were found for any of the construction periods, or the maintenance period, in the posterior hippocampus.

**Figure 3.**
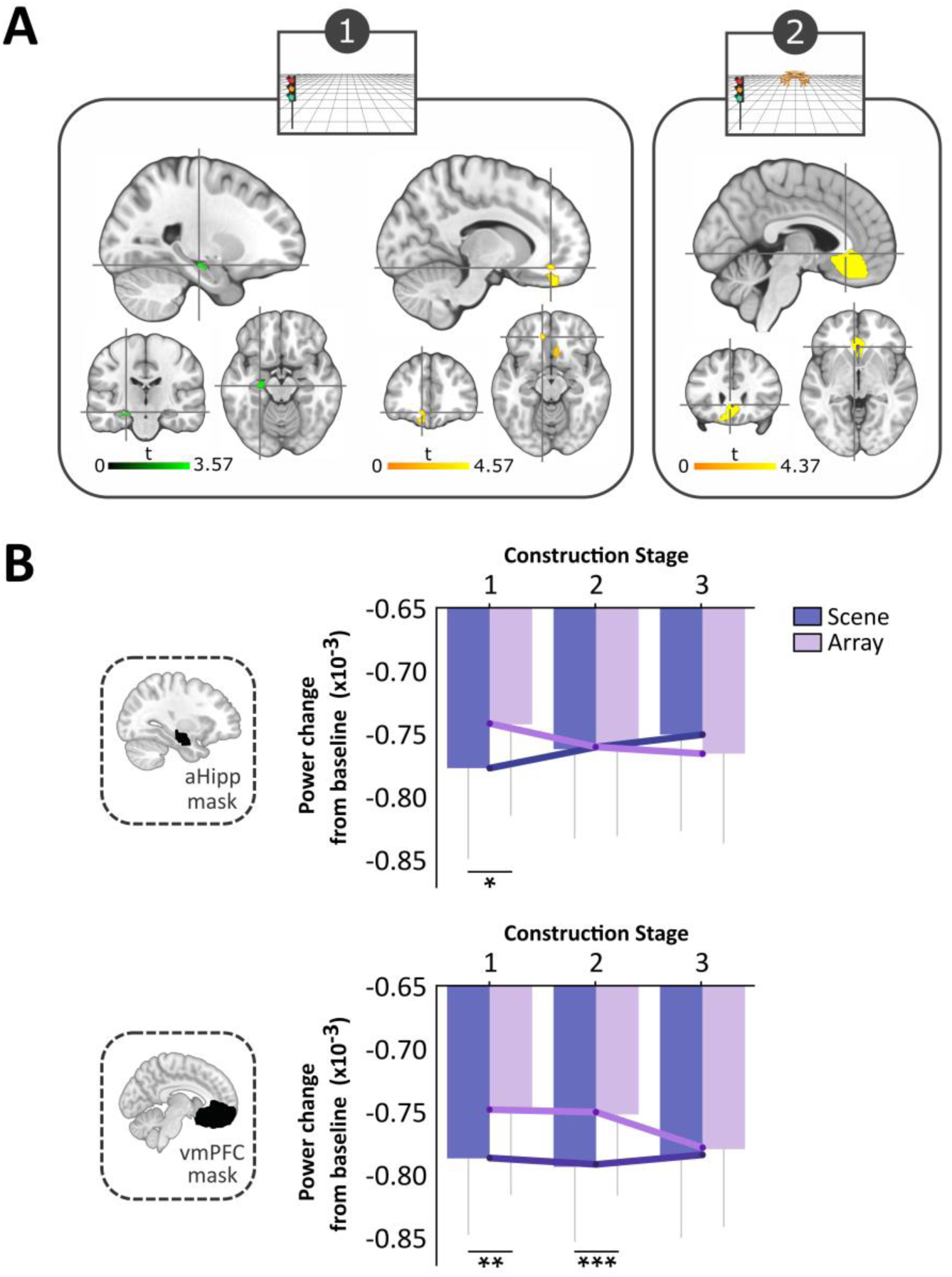
MEG results. (A) Significant theta power changes within our two regions of interest, the anterior hippocampus (aHipp, shown in green) and ventromedial prefrontal cortex (vmPFC, shown in yellow) during scene relative to array construction (FWE p < 0.05, corrected for the ROI). Slices are centred on the overall peaks (crosshairs) in the left anterior hippocampus (x, y, z = −24, −20, −16) and left vmPFC (−12, 40, −14) during the first construction stage, and the vmPFC (−4, 26, −4) during the second period of construction. (B) Task × construction stage interactions for the change in theta power relative to baseline in the anterior hippocampus and vmPFC. For the hippocampus, power was significantly attenuated for the scene relative to the array condition during the first stage of construction. For the vmPFC, power was significantly attenuated for the scene compared to the array task for both the first and second construction periods. *p = 0.014, **p = 0.008, ***p = 0.001. Error bars indicate ± 1 SEM.

To investigate whether there were differences between scene and array tasks in frequencies other than theta, we performed the same analyses for alpha (9-12 Hz) and gamma (30-100 Hz) bands. No significant changes were evident in the anterior (or posterior) hippocampus or vmPFC.

### Theta Power Changes Across Construction Stages

Having observed between-task differences when each construction stage was considered separately, we next analysed the data in a different way to examine how theta activity changed across the construction stages. For both scene and array conditions, theta power was attenuated across all stages relative to baseline (Figure 3B), which we regard as reflecting increased task-related neural activity. An ANOVA assessing the effects of task (scene, array) and construction stage (first, second, third) on anterior hippocampal activity revealed no significant effect of task (*F*_(1, 19)_ = 0.694, *p* = 0.415) or construction stage (*F*(_2, 38)_ = 0.036, *p* = 0.965), but a significant task × construction stage interaction (*F*_(2, 38)_ = 5.762, *p* = 0.007; Figure 3B). This was driven by a significantly greater theta power decrease during the first construction stage of the scene task (*t*_(19)_ = −2.713, *p* = 0.014), indicating increased involvement of the anterior hippocampus during this scene stage, with engagement then decreasing over the subsequent two construction periods.

For the vmPFC, the ANOVA revealed a significant main effect of task (*F*_(1, 19)_ = 17.252, *p* = 0.0005; Figure 3B), but not of construction stage (*F*_(1.488, 28.272)_ = 0.894, *p* = 0.392), and there was no significant interaction (*F*_(2, 38)_ = 2.586, *p* = 0.089). The task effect was driven by larger theta power decreases during the scene relative to the array task during both the first (*t*_(19)_ = −2.959, *p* = 0.008) and second (*t*_(19)_ = −3.723, *p* = 0.001) construction stages, indicating the greatest vmPFC involvement during these periods of the scene task.

### Effective Connectivity During Scene Construction Stages

The source localisation analyses described above established the peak locations of theta power changes in the anterior hippocampus and vmPFC during scene compared to array construction. We next examined the direction of information flow between these regions during each task using DCM for cross spectral densities using the left hemisphere peaks in the anterior hippocampus and vmPFC identified during the first construction stage (Figure 3A). We proposed three connectivity models based on established anatomical projections, representing the hypotheses for the hierarchical architecture of the hippocampus and vmPFC (Figure 4). The first model predicted hippocampal activity drove that of the vmPFC. Hippocampal projections via the fornix terminate in the middle layers of the ventral portion of the vmPFC (Aggleton, Wright, Rosene, & Saunders, 2015), so this connection was classified as ‘forward’. The second model specified the vmPFC as the influencing region. Connections from the vmPFC to the hippocampus are via the entorhinal cortex, where vmPFC inputs arrive at all cell populations (Rempl-Clower and Barbas, 2000), so were classified as ‘lateral’ connections. The third model predicted that the hippocampus and vmPFC drove each other through reciprocal connections (mutual entrainment), so both ‘forward’ and ‘lateral’ connections were specified. We then fit the models to each of the three 3000 ms scene and array construction stages. The approach adopted here to classifying DCM connections is also in line with previous work modelling hippocampal and vmPFC interactions in MEG (Barry, Barnes, Clark, & Maguire, 2019). As an extension to this previous DCM framework, we not only compared alternative unidirectional ‘master-slave’ relationships, but also directly compared these models with bidirectional mutual entrainment, where neither region exerted a stronger influence over the other.

**Figure 4.**
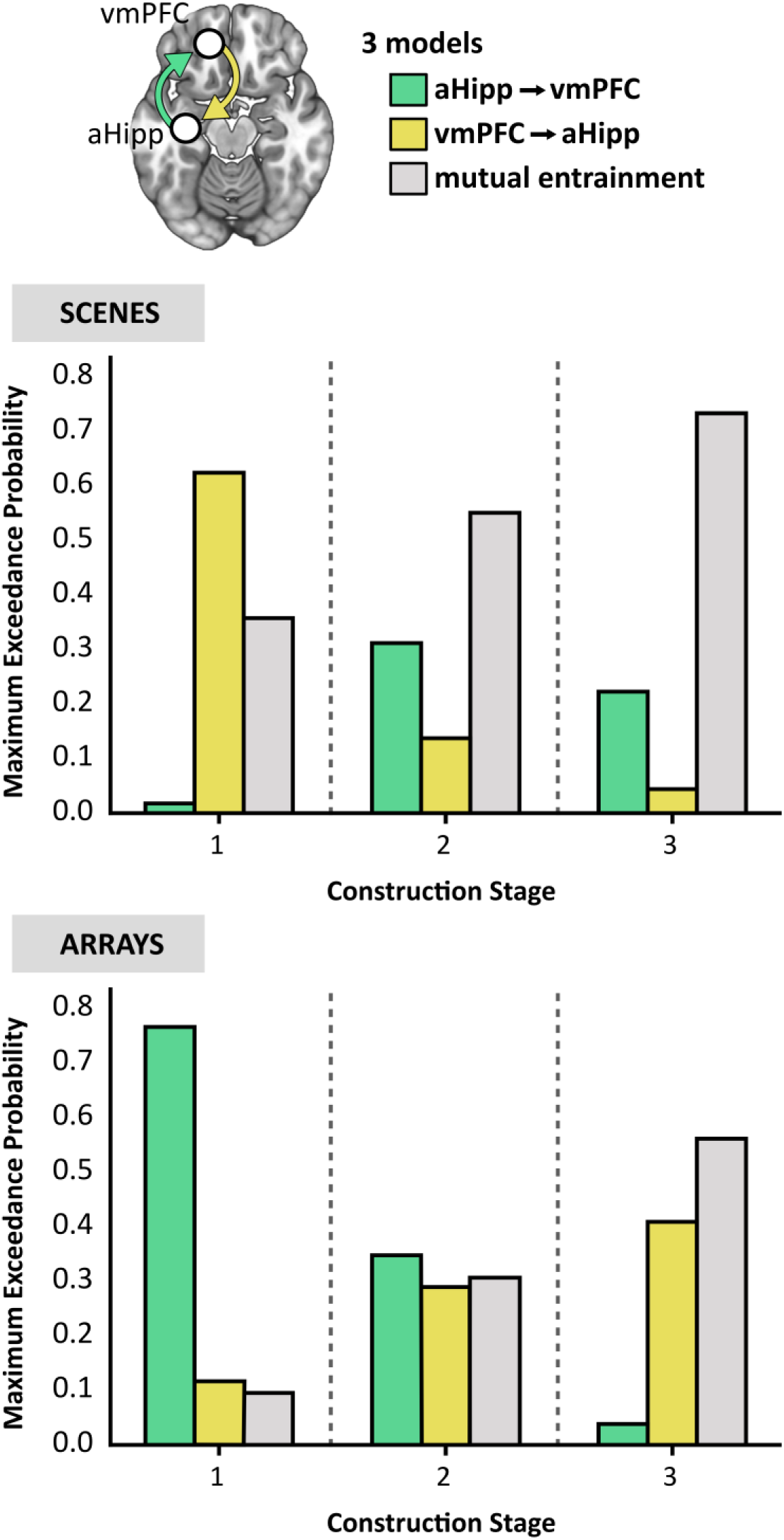
Connectivity between the anterior hippocampus (aHipp) and ventromedial prefrontal cortex (vmPFC) during step-by-step scene and array construction. Top panel, the three models of effective connectivity. Middle panel, the results of Bayesian model selection revealed the model with the strongest evidence during scene construction stage 1 was the vmPFC driving anterior hippocampus, while for the latter two stages, there was stronger evidence for the anterior hippocampus and vmPFC driving each other through reciprocal connections (mutual entrainment). Bottom panel, for array construction, the winning model during construction stage 1 was the anterior hippocampus driving the vmPFC, for the second stage there was no clear evidence for any model, and for the third stage there was stronger evidence for mutual entrainment.

The models were compared using random effects Bayesian model selection (BMS) to determine the winning model at each construction period at the group level (Figure 4). During the initial construction period for scenes, the model most likely to explain the data across all participants was the vmPFC driving anterior hippocampal activity, with an exceedance probability of 62.45%, almost double that for the mutual entrainment model (exceedance probability = 35.78%). Furthermore, there was very little evidence for the opposing model of the anterior hippocampus driving vmPFC (exceedance probability = 1.77%). During the second scene construction stage, the mutual entrainment model was the most probable, with an exceedance probability of 55.08%, compared to the hippocampus-driven model (exceedance probability = 31.18%) and the vmPFC-driven model (exceedance probability = 13.74%). During the third scene construction stage, the winning model was again the mutual entrainment model (exceedance probability = 73.32%) compared to the hippocampus-driven model (exceedance probability = 22.28%) and the vmPFC-driven model (exceedance probability = 4.40%).

DCM of array construction revealed a different pattern of effective connectivity between the two ROIs. At the first stage of array construction, the model that best fitted the data was the anterior hippocampus driving vmPFC activity, with an exceedance probability of 76.74%. There was little evidence for either the vmPFC driving the hippocampus (exceedance probability = 12.06%) or mutual entrainment (exceedance probability = 11.20%). During the second array construction stage, there was no clear winning model (exceedance probabilities = hippocampus-driven 38.54%; vmPFC-driven 30.38%; mutual entrainment 31.08%). During the third array construction stage, the mutual entrainment model was the most probable (exceedance probability = 56.10%), followed closely by the vmPFC-driven model (exceedance probability = 40.75%), and least likely was the hippocampus-driven model (exceedance probability = 3.15%).

## DISCUSSION

In this study we focused on the anterior hippocampus and vmPFC and leveraged the fine temporal resolution of MEG to investigate how their neural dynamics were associated with the step-by-step mental construction of scene and non-scene array imagery. Across three successive construction stages, we observed scene-related attenuation of theta power in the anterior hippocampus during the initial stage, and in the vmPFC during the first two stages, relative to array construction. We also found that the scene-specific anterior hippocampal theta activity during the first phase of construction was driven by the vmPFC, with mutual entrainment between the two brain regions thereafter. By contrast, array construction was associated with the anterior hippocampus driving vmPFC activity during the first construction stage.

The power changes we observed were, as predicted, specific to theta, aligning with the long association between theta and hippocampal function (reviewed in Colgin, 2016; Kerakas, 2020), and the previous MEG study of immediate full scene construction of Barry, Barnes, Clark, & Maguire (2019). Our findings of significantly greater attenuation in theta power for the first (anterior hippocampus, vmPFC) and second (vmPFC) stages of scene relative to array construction, which we consider to reflect increased task-related neural activity, also accords with the extant literature (e.g. Fellner et al., 2016; Fellner et al., 2019; Solomon et al., 2019; Barry, Barnes, Clark, & Maguire, 2019).

Our primary interest was in whether increased engagement of the anterior hippocampus and vmPFC would be specific to scene construction. As predicted, both brain regions were preferentially recruited during the mental construction of scenes compared to arrays particularly during the early stages. Our results converge with the previous fMRI findings of Dalton, Zeidman, McCormick, & Maguire (2018) and the MEG findings of Barry, Barnes, Clark, & Maguire (2019), and extend them in several ways. For example, in the Barry, Barnes, Clark, & Maguire (2019) study, participants imagined a full scene within three seconds, and this was compared to a low level baseline (imagining single objects). Dalton, Zeidman, McCormick, & Maguire (2018) compared scenes and arrays but could not examine the three construction stages separately given the slow nature of the haemodynamic response. By contrast, in the current study we examined the gradual step-by-step construction of scenes and showed that even at the earliest stage, with one object and a simple 3D space, this was sufficient to engage the anterior hippocampus and vmPFC, and more so than the first stage of array construction. Hence we could drill down further than before into how a scene develops, with a much tighter comparison in the form of the arrays, and in doing so show that a scene is ‘set’ early on.

We considered the scene and array tasks to be very well matched in terms of the requirement for constructive and associative processing. However, it could be argued that the 3D space in the scenes somehow required more relational processing than the 2D space in the arrays, although it is unclear how the experience of depth, inherent to scenes, would translate into additional associative processing. Of note, when the imagined 3D and 2D spaces alone (without objects) were examined during fMRI by Dalton, Zeidman, McCormick, & Maguire (2018), neither was associated with increased hippocampal activity (see also Zeidman, Mullally, Schwarzkopf, & Maguire, 2012). Moreover, proponents of the relational theory are clear that mental operations involving 3D space, such as navigation, engage the hippocampus because they are an example of associative processing, and that non-spatial stimuli involving associations should also engage the hippocampus. Consequently, and given the close similarity of the stimuli in our scene and array tasks, we believe that, a priori, relational theorists would have predicted similar hippocampal involvement for the scenes and arrays, which we did not find.

Nevertheless, if the scene stimuli involved more associative processing, how might we detect this in our data? We may have expected to observe a difference between scenes and arrays in terms of eye movements, yet none were evident. If more relational processing was required for scenes then we might have predicted differential performance on the surprise post-scan item memory test, but none was found. Of note, when Dalton, Zeidman, McCormick, & Maguire (2018) examined associative memory for the same stimuli in a surprise post-fMRI scan memory test, there was also no difference between scenes and arrays (in fact, both were at chance, as this is a very challenging task). Given the simplicity of our stimuli, three objects and a space, and their similarity on these measures, we believe that a parsimonious interpretation of our paradigm is that the two conditions were comparable in terms of associative processing.

Why might scene construction be particularly engaging for the anterior hippocampus and vmPFC? Scene imagery seems to be important for, and is frequently deployed in the service of, hippocampal-dependent tasks including episodic memory, imagining the future and spatial navigation. For example, using a large sample (n = 217) of participants and multiple cognitive tests with a wide spread of individual differences in performance, Clark et al. (2019) reported that the ability to construct scene imagery explained the relationships between episodic memory, imagining the future and spatial navigation task performance. The prominence of scene imagery was further emphasised in another study involving the same sample, where the explicit strategies used to perform episodic memory recall, future thinking and spatial navigation tasks was assessed (Clark, Monk, & Maguire, 2020). In each case, the use of scene visual imagery strategies was significantly higher than for all other types of strategies (see also Andrews-Hanna, Reidler, Sepulcre, Poulin, & Buckner, 2010). The influence of scene imagery across cognition is perhaps not surprising, given that it mirrors how we perceive the world.

Our findings also provide insights into the interactions between the anterior hippocampus and vmPFC during scene construction. McCormick, Ciaramelli, De Luca, & Maguire (2018) and Ciaramelli, De Luca, Monk, McCormick, & Maguire (2019) proposed that the vmPFC initiates the activation of schematic and other knowledge in neocortex that is relevant for a scene, whilst inhibiting elements that are not relevant. This information is conveyed to the hippocampus which constructs the scene. vmPFC then engages in iterations via feedback loops with neocortex and hippocampus to mediate the schema-based retrieval and monitoring necessary to flesh-out the scene. That the vmPFC drove activity in the anterior hippocampus during the first scene construction phase supports the perspective that the vmPFC instigates this process. After this initial scene-setting, hippocampal and vmPFC activity was mutually entrained, perhaps reflecting iterative feedback loops between them as each new object was integrated to form a fully realised scene.

The activation of schemas by vmPFC may provide an initial rudimentary template into which a scene may be constructed in collaboration with the hippocampus. Although our scene stimuli were composed of object triplets that were not contextually associated (van Kesteren, Ruiter, Fernández, & Henson, 2012), there is evidence for schema-guided construction of novel representations driven by the vmPFC (Garrido, Barnes, Kumaran, Maguire, & Dolan, 2015). Furthermore, the observed tonic response of the vmPFC throughout the first two construction stages in our study may reflect the prediction and preparatory processing necessary for the subsequent integration of scene elements (Ciaramelli, De Luca, Monk, McCormick, & Maguire, 2019; McCormick, Ciaramelli, De Luca, & Maguire, 2018).

A very different result was evident when connectivity between our two ROIs was examined during the array condition. For the first stage of construction, the anterior hippocampus drove vmPFC activity. Why we observe this direction of connectivity for the arrays is not clear. We speculate that during array construction, with the vmPFC much less engaged compared to the scenes during the first two construction stages, perhaps the hippocampus was ‘reaching out’ for guidance from the vmPFC. Interestingly, at the third stage, the vmPFC was equally engaged for arrays and scenes, and the DCM analysis showed mutual entrainment to be the most probable explanatory model. This may indicate that the vmPFC stepped in to provide some input. Of course, this is conjecture, and future studies are required to explore this finding further.

In conclusion, the hippocampus and vmPFC are particularly implicated in supporting cognitive functions such as episodic memory, imagining future events and spatial navigation. While it is likely these two brain areas participate in associative and constructive processing more generally (Dalton, Zeidman, McCormick, & Maguire, 2018), here we have shown that their neural activity seems to be especially tuned to the earliest stage of building scene representations. In future studies it will be important to elucidate the contributions of other brain regions, including posterior cortical areas (Dalton & Maguire, 2017), to this process and to characterise putative anatomical pathways that facilitate hippocampal-vmPFC interactions in the human brain. Moreover, if, as some have argued, scenes are the smallest functional building blocks of extended mental events (Ciaramelli, De Luca, Monk, McCormick, & Maguire, 2019), we also need to understand how individual scenes are combined to construct the seamless dynamic unfolding episodes that constitute the memories of our past experiences.

## Acknowledgements

This work was supported by a Wellcome Principal Research Fellowship to E.A.M. (210567/Z/18/Z) and the Centre by a Centre Award from the Wellcome Trust (203147/Z/16/Z). We are grateful to the attendees of the MEG meetings at the Wellcome Centre for Human Neuroimaging at UCL for their advice. Thanks also to Daniel Bates, David Bradbury and Eric Featherstone for technical support. The authors declare no competing financial interests and no conflicts of interest.

